# Programmable DNA-Silk fibroin hydrogel scaffold with temporal mechanical reinforcement for mesenchymal stem cells differentiation and accelerated wound healing in a full-thickness skin defect mice model

**DOI:** 10.1101/2025.11.06.686972

**Authors:** Nihal Singh, Bhagyesh Parmar, Aneri Joshi, Devanshi Gajjar, Vivek Kashyap, Raghu Solanki, Ankur Singh, Seshadri Sriram, Dhiraj Bhatia

## Abstract

DNA-based hydrogels have attracted significant attention in biomedical applications due to their programmability, biocompatibility, and tunable functionalities. However, their clinical translation is hindered by high synthesis costs, limited mechanical strength, poor stability, and complex synthesis protocols. In this work, we present a novel DNA-silk fibroin hybrid hydrogel that overcomes these limitations by combining the unique biofunctionality of DNA with the superior mechanical and chemical properties of silk fibroin. Salmon-derived DNA, an abundant and low-cost source, was incorporated with silk fibroin to form hybrid networks via a simple, scalable process. Silk fibroin served as a robust scaffold, enhancing mechanical strength, controlling degradability, and improving structural stability, while the entangled DNA-silk network reduced rapid DNA degradation. The resulting hybrid hydrogel demonstrated temporal mechanical reinforcement and prolonged stability, with in vitro studies demonstrating that the developed hybrid hydrogel enhanced the chondrogenic and osteogenic differentiation of Infrapatellar Fat Pad Mesenchymal Stem Cells (IFP-MSCs). Additionally, hemostatic and in vivo studies demonstrated the potential of DNA-silk fibroin hydrogel as a promising biomaterial for rapid hemostasis and accelerated wound healing. Overall, the developed DNA-silk fibroin hybrid hydrogel offers strong potential for mesenchymal tissue engineering, hemostatic adjuvant and wound healing.

## 1. Introduction

DNA, a biopolymer selected by nature itself to be the carrier of genetic information, is now considered an important structural material owing to its high modification capacity at both the genomic and molecular level^1^. Utilizing the predictable Watson-Crick base pairing, DNA-based soft materials with complex architecture, inherent programmability, biocompatibility, and stimuli responsiveness have been developed^2,3^. Today, DNA is utilized to develop functional biomaterials spanning across the nanoscale, microscale, and bulk scales^4,5^. At the nanoscale, DNA nanotechnology leverages DNA’s nanoscopic scale and programmable nature to develop 2D and 3D nanomaterials and nanodevices with controllable properties^6–8^. In contrast, at the microscale, DNA programmable self-assembly is utilized to fabricate nanofibers, nanotubes, and nanosheets^9–12^. At the bulk or macroscopic scale, DNA is primarily utilized as a hydrogel utilizing the hydration of negatively charged sugar-phosphate backbones, creating hydrogel-like structures^13,14^. Today, DNA-based hydrogels have attracted significant attention owing to their biocompatibility, biodegradability, programmable molecular recognition, catalytic activities, and therapeutic potential. Additionally, temperature, pH, ionic strength, and solvent composition responsive DNA hydrogels have also been developed, offering advanced functionalities for biomedical applications, including controlled drug delivery, targeted gene therapy, cancer therapy, biosensing, protein production, 3D cell culture, wound healing, and tissue engineering^15^. However, despite significant advancement and broad application potential, DNA-based hydrogels suffer from a number of significant limitations that impede their widespread application in biomedical, biosensing, environmental, and tissue engineering fields.

In recent years, significant research has utilized DNA hydrogel for a wide range of applications; however, the transition of DNA hydrogel from proof-of-concept laboratory systems to robust, scalable platforms for clinical and translation use has been hindered by several intrinsic and extrinsic limitations. One of the primary limitations of DNA-based hydrogels is the high cost of DNA synthesis^16^. Hydrogels inherently require a large amount of hydrophilic polymer to form a network that can absorb and retain a large amount of water. Thus, DNA-based hydrogel at the micro and macroscale often requires DNA in milligrams to perform as a hydrogel. This significantly increases the synthesis cost, hindering scale-up to the translation stage. The high synthesis cost is further exacerbated when production of DNA with precise sequences and modifications is required, which is often the case with self-assembled DNA hydrogel^17^. Moreover, techniques such as rolling circle amplification (RCA), which are employed to generate long DNA strands, can simplify design and reduce the cost per strand; however, these enzymatic methods tend to be time-consuming and introduce additional challenges related to purification and batch-to-batch variability^18,19^. Another major limitation of DNA hydrogels is their poor mechanical strength and stability, which becomes highly apparent when used for applications such as wound healing and tissue engineering^20,21^. The intrinsic properties of DNA, such as high persistence length and large mesh size, result in a hydrogel with low stiffness and poor mechanical robustness. This significantly limits the potential of DNA hydrogels for applications that require appropriate mechanical properties, such as tissue engineering. However, the stability of DNA hydrogels under physiological conditions is of the most significant concern. Cellular and enzymatic degradation of DNA hydrogels in a biological setting often results in rapid degradation and loss of function, with an operational lifespan of just a few hours^22^. Even DNA hydrogels with high crosslinking density degrade within a few days in biological fluids. Furthermore, cell culture studies are also not feasible, as DNA hydrogels rapidly degrade in cell culture media supplemented with serum. Such instability is particularly problematic when the hydrogel is intended for prolonged therapeutic or tissue engineering applications, where controlled degradation kinetics and sustained mechanical integrity are essential^23^. Hence, while the unique properties of DNA-based hydrogels offer a new paradigm for smart material design, their present limitations in synthesis cost, mechanical robustness, and biological stability pose significant hurdles that must be addressed before DNA-based hydrogels can achieve widespread practical application.

To address the above limitations of DNA-based hydrogels, one promising approach is the development of hybrid hydrogels, where DNA is used in conjunction with other polymers that can provide mechanical strength, chemical versatility, and improved stability^2,24,25^. Such systems may help mitigate the high cost, limited stability, poor mechanical strength, and complex synthesis requirements associated with pure DNA hydrogels while retaining the inherent advantages of DNA. Towards this aim, we successfully developed a DNA-Silk fibroin hybrid hydrogel with superior mechanical properties, higher stability, easy synthesis, and low synthesis cost. To reduce the cost associated with DNA synthesis, we used DNA from salmon, which is abundantly available, making the hydrogel cost-efficient, enabling scale-up for clinical applications. We used silk fibroin as a scaffold for hydrogel, as silk fibroin is a natural polymer with exceptional biocompatibility, controllable degradability, and superior mechanical properties^26,27^. Utilisation of silk fibroin as a scaffold imparted DNA-silk hydrogel with superior mechanical properties, enabling temporal mechanical reinforcement, and simultaneously, the entangled network formed from DNA and silk crosslinking prevented rapid degradation of DNA. Additionally, to investigate the suitability of the developed DNA-Silk fibroin hydrogel for biological applications, we investigated the potential of DNA-Silk fibroin hydrogel as a scaffold for mesenchymal stem cell differentiation for cartilage and bone tissue engineering applications. Also, as DNA is an essential component of the neutrophil extracellular traps (NETs) matrix, which is involved in blood clotting and coagulation, we investigated the potential of DNA-Silk hydrogel as a hemostatic adjuvant with application in wound healing. Overall, we propose a novel DNA-silk fibroin hybrid biomaterial that overcomes the major limitations associated with pure DNA-based materials and provides a possible strategy for the utilisation of DNA-based material for applications such as mesenchymal cell differentiation, rapid homeostasis, wound healing and tissue engineering.

## 2. Results and Discussion

### 2.1 Synthesis and morphological characterisation of DNA-Silk fibroin hydrogel

The synthesis of DNA-Silk fibroin hydrogels was achieved using a simple one-pot assembly process **(Figure 1A)**. First, the silk fibroin solution was extracted from the B. *mori* cocoons, and the DNA from the salmon testes was dissolved in nuclease-free water. The DNA was mixed with silk fibroin solution in different ratios to form a hydrogel pre-gel solution **(Table 2)**. Pure DNA solution without silk fibroin was taken as the DNA control (1.5G), and pure silk fibroin solution without DNA was taken as the silk fibroin control (2S and 4S). First, crosslinking and a supramolecular network formation between the DNA strands was achieved by heating the hydrogel pre-gel solution to 93°C. This exposure of DNA to high temperatures breaks hydrogen bonds between base pairs, rendering the DNA single stranded. The resulting single-stranded DNA network was rapidly cooled to 37°C to promote non-specific base pairing, leading to random crosslinking. This non-specific crosslinking of DNA strands resulted in rapid hydrogel formation within 15 min with an entangled silk fibroin polymer network. However, the transparent solution of the initial DNA-Silk hydrogel indicated that the silk fibroin chains in the hydrogel are still in random coil form and have not transitioned to β-sheet structure **(Figure 1B)**. However, after 48 h, the DNA-Silk hydrogel started to become translucent, indicating the transition of silk fibroin chains to β-sheet conformation^28,29^. After 72 h, the DNA-Silk fibroin hydrogel became highly translucent, indicating a significant transition of silk random chains to β-sheet conformation. The transition of silk fibroin from a random coil to a β-sheet conformation, which typically takes a month, can be attributed to the exposure of silk fibroin to high temperature during synthesis and subsequent crosslinking of a DNA network, which constrains and brings silk fibroin chains together, resulting in a β-sheet transition. To confirm the presence of β-sheet formation in DNA-Silk fibroin hydrogel, we performed FTIR analysis in the range of 1600 to 1700 cm^-1^. A characteristic peak at 1633 cm^-1^, which is associated with β-sheet, was observed, indicating the transition of silk fibroin to β-sheet in DNA-Silk fibroin hydrogel **(Figure 1C)**. To further observe the morphology and structure of DNA-Silk fibroin hydrogel, SEM micrograph images of the hydrogels maintained at 37° C for 5 days were taken **(Figure 1D)**. As observed, both 1.5D2S and 1.5D4S hydrogels exhibited a highly porous architecture with 1.5D2S and 1.5D4S hydrogels having a pore size of 87.66 ± 20.85 µm and 60.13 ± 16.22 µm, respectively **(Figure 1E)**. These observations suggest that as the concentration of silk fibroin within the hydrogel increases, the pore size of the hydrogel decreases. In contrast, the pure DNA hydrogel demonstrated an interconnected fibrous network at the surface with a dense film architecture with very few pores at the bottom. This interconnected fibrous network can be due to the single DNA strands crosslinking during synthesis, validating that the non-specific crosslinking of DNA strands results in a supramolecular network formation. However, this interconnected fibrous network architecture was not observed in DNA-Silk fibroin hydrogel; instead, a highly porous architecture was observed in DNA-Silk fibroin hydrogel, suggesting that silk fibroin acts as a molecular scaffold and alters the architecture of DNA hydrogel from fibrous to porous.

**Figure 1:**
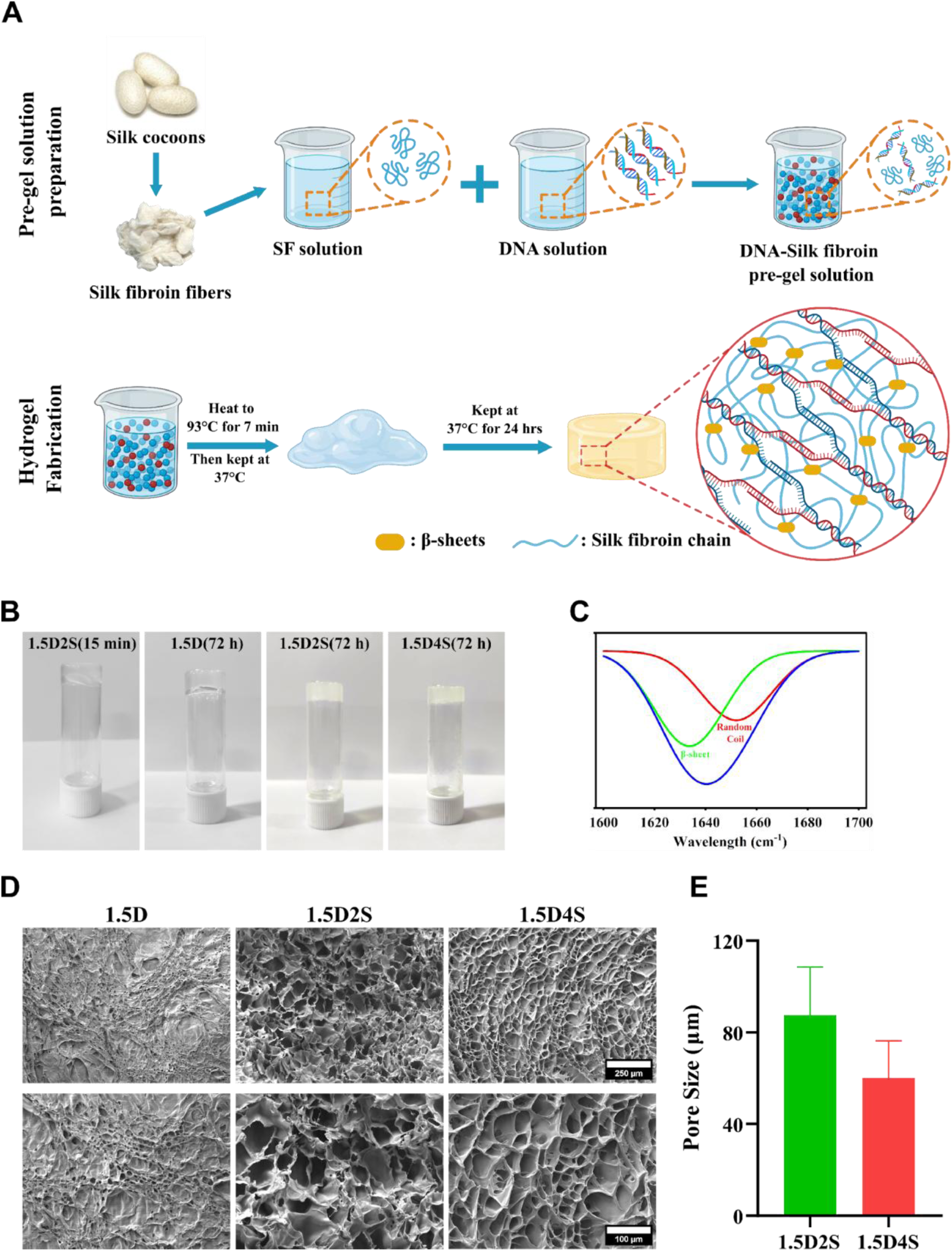
Synthesis and characterization of DNA-Silk fibroin hydrogel. (A) Schematic of the synthesis of DNA-Silk fibroin hydrogel. (B) Vial inversion test for the developed hydrogels. (C) Deconvoluted FTIR spectrum of DNA-Silk fibroin hydrogel. (D) SEM micrographs images of developed hydrogel systems. (E) Quantification analysis for the pore size of DNA-Silk fibroin hydrogels using ImageJ. Data presented as mean ± SD.

### 2.2 Chemical and mechanical characterisation of DNA-Silk fibroin hydrogel

To further understand the mechanism of DNA-Silk fibroin hydrogel formation at the molecular level, we performed FTIR analysis of DNA, silk fibroin, and DNA-Silk fibroin hydrogels. A characteristic FTIR spectrum for DNA hydrogel was observed with FTIR peaks at 1050 cm^-1^ and 1220 cm^-1^ corresponding to the DNA phosphate backbone (POC/PO₂⁻) vibrations, respectively **(Figure 2A)**. Additionally, the broad band at 3200-3400 cm⁻¹ along with the peaks at 1650 cm⁻¹ corresponding to OH and NH stretching of DNA bases and sugar and C=O and C=N stretching in nucleobases, respectively, confirmed the presence of DNA components: phosphate, sugar, and bases. Similarly, a characteristic FTIR spectrum of a protein sample was observed in the case of silk fibroin with characteristic peaks at 1625 cm^-1^, 1527 cm^-1^, and 1230 cm^-1^ corresponding to amide I, amide II, and amide III observed in the protein sample, respectively. However, in DNA-Silk fibroin hydrogel, which is a combination of both silk and DNA, several notable changes can be observed **(Table 1)**. First, combined peaks of both DNA and silk fibroin were observed in the DNA-Silk fibroin hydrogel, indicating that the hydrogel contains a completely intertwined and homogeneous mixture of both DNA and silk fibroin. Further, a broadening and shift in OH/NH Stretch around 3280 cm⁻¹ was observed, indicating a possible stronger hydrogen bonding interaction between DNA’s phosphate groups and silk’s amide NH groups. Similarly, a slightly weaker and broadened peak around the phosphate stretch (1000-1250 cm⁻¹) was observed in DNA-Silk fibroin hydrogel, which can be indicative of possible electrostatic interactions of DNA with Silk fibroin. Additionally, changes in amide I (weakened, slightly shifted) and amide II (reduced intensity) peaks point to partial disruption or conformational change in Silk’s fibroin β-sheet structure due to interactions with DNA. Overall, the FTIR data validate the successful formation of a DNA-Silk fibroin hydrogel and provide deep insight with strong evidence of non-covalent interactions such as hydrogen bonding and electrostatic attraction. It further suggests that these interactions resulted in partial structural reorganisation, which resulted in a stable hybrid material formed through the non-covalent network.

**Figure 2:**
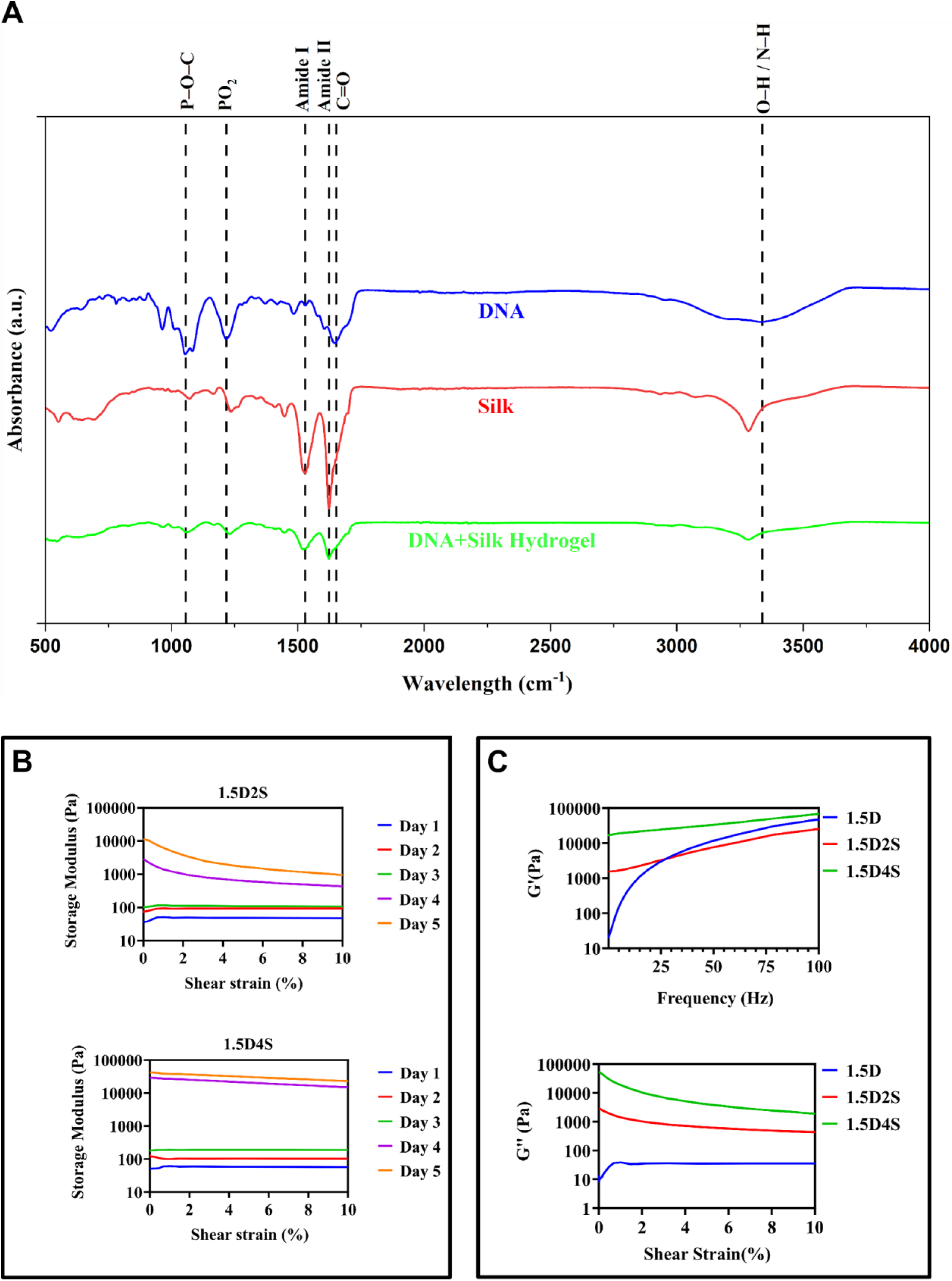
Chemical and mechanical characterisation of DNA-Silk fibroin hydrogel. (A) FTIR spectrum of developed hydrogels. The dashed line represents the corresponding functional group associated with the wavenumber. (B) Time-dependent analysis of the rheological properties of the developed DNA-Silk fibroin hydrogel. (C) Comparison of rheological properties of pure DNA hydrogel (1.5D) and DNA-Silk fibroin hydrogel (1.5D2S and 1.5D4S) utilising frequency sweep and amplitude sweep of the hydrogels.

**Table 1:** Summary of FTIR spectral changes and interpretation for DNA-Silk fibroin hydrogel.

| Spectral Feature | Change in Hydrogel | Interpretation |
| --- | --- | --- |
| O–H/N–H stretch<br>($\sim 3280\text{ cm}^{-1}$) | Broadened, shifted | Stronger hydrogen bonding between DNA and silk |
| Amide I ($\sim 1625\text{ cm}^{-1}$) | Weakened, slightly shifted | Disruption of silk β-sheet structure |
| Amide II ($\sim 1527\text{ cm}^{-1}$) | Reduced intensity | Silk secondary structure modification |
| Phosphate stretch<br>($1000\text{--}1250\text{ cm}^{-1}$) | Broadened, slightly weaker | Electrostatic/H-bonding interactions |
| Overall peak sharpness | Decreased | Blending into a new hydrogel network |

**Table 2:**
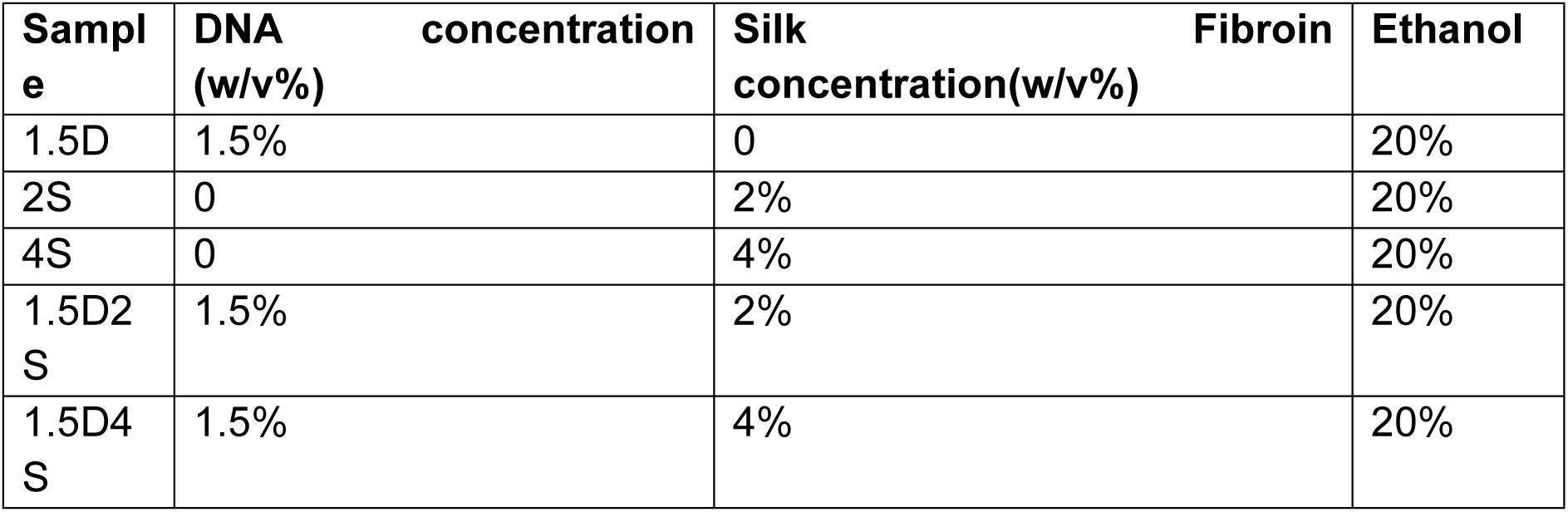
Different weight percentage ratios of DNA and Silk fibroin for the preparation of DNA-Silk hybrid hydrogel.

To further understand the effect of the incorporation of silk fibroin on the mechanical properties of DNA-Silk fibroin hydrogel, rheological studies were performed. As the transition of silk fibroin from random coil to β-sheet confirmation in DNA-Silk fibroin hydrogel was not immediate and occurred over several days, we wanted to investigate whether this gradual transition also affects the mechanical properties of the hydrogel. Hence, the mechanical properties of DNA-Silk fibroin hydrogels were evaluated every 24 h. We observed a continuous increase in the mechanical strength of hydrogels 1.5D2S and 1.5D4S over the period of 5 days, after which no further increase in mechanical strength was observed **(Figure 2B)**. Both hydrogels exhibited a marked increase in storage modulus (G’), suggesting enhanced crosslinking density and increased structural reinforcement of the DNA-Silk fibroin hydrogel is due to the incorporation of silk fibroin, which contributes to the maturation and mechanical robustness of the hydrogel matrix. Further, the viscoelastic behaviour of pure DNA hydrogel and DNA-Silk hydrogel was evaluated through frequency and amplitude sweep analysis. We observed that both 1.5D2S and 1.5D4S hydrogels demonstrated significantly higher storage modulus in comparison to the 1.5D hydrogel, demonstrating that the incorporation of silk fibroin in the DNA network can significantly increase the mechanical strength of the pure DNA hydrogel **(Figure 2C)**. Also, 1.5D4S hydrogel demonstrated higher mechanical reinforcement in comparison to 1.5D2S hydrogel, indicating that the increased mechanical reinforcement is silk fibroin concentration dependent and as the concentration of silk fibroin increases, the rate as well as the strength of mechanical reinforcement increases. Overall, these results demonstrate that the incorporation of silk fibroin in pure DNA hydrogel can overcome the limitation of poor mechanical strength associated with DNA hydrogel. Additionally, due to the gradual β-sheet transition in DNA-Silk fibroin hydrogel, a highly dynamic hydrogel with dynamic stiffening was developed. These results further extend the potential application of the DNA-Silk hydrogel, such as tissue engineering, where the DNA-Silk hydrogel can provide time-dependent mechanical support, gradually mimicking the developing strength of regenerating tissues such as cartilage, muscle, or vascular grafts. Similarly, the developed hydrogel can be used in drug delivery systems for the dynamic drug release of therapeutic drugs at the local site with decreased diffusion rate as the crosslinking density of the hydrogel increases^30^. Overall, the developed hydrogels with dynamic mechanical reinforcement can be used for various biological applications, making them a potential candidate for next-generation biomaterials.

### 2.3 Degradation behaviour of DNA-Silk fibroin hydrogel

One of the significant limitations of pure DNA-based hydrogels is their limited stability in a biophysiological environment. This limited stability of the DNA hydrogel, both in vivo as well as in cell culture medium, can be attributed to DNA strand cleavage by nucleases as well as to salt-induced DNA expansion^22,23^. DNA hydrogels with a lower DNA concentration are especially susceptible to cleavage by DNase I due to their limited crosslinked density, providing easy access to DNase I for cleavage, leading to gradual degradation and complete dissolution of the DNA hydrogel. Attempts to increase crosslinking density by increasing the concentration of DNA have shown higher resistance to degradation, but significantly increase synthesis cost due to the amount of DNA required. Similarly, the presence of high salts also triggers destabilisation of the condensed hydrogels, which precedes enzyme-induced degradation^31^. The onset of degradation correlates with high salt-induced DNA hydrogel expansion, whereby monovalent cations like Na+ and K+ destabilise the condensed structure, exposing DNA strands to enzymatic cleavage. Furthermore, most cell culture medium also contain serums such as fetal bovine serum, which contains various nucleases, resulting in hydrogel breakdown. Thus, physiological ionic compositions and nucleases jointly contribute to hydrogel instability and fast degradation in in vivo and cell culture conditions, significantly limiting the application of DNA-based hydrogels^32^. Most studies report the stability of DNA hydrogel in water or PBS, where, due to the absence of the DNase enzyme, a correct estimation of DNA hydrogel stability cannot be estimated.

Therefore, we studied the degradation profile of DNA-Silk fibroin hydrogel in both PBS and cell culture medium. We aimed to investigate whether DNA-silk fibroin can resist DNA degradation and thereby extend the stability of the hydrogel, particularly for applications requiring a slow degradation rate, such as wound healing, drug delivery, and tissue engineering. To explore the stability of DNA-Silk fibroin hydrogel, the hydrogels were incubated with PBS and DMEM cell culture media. At specific timepoints, the PBS and DMEM cell culture media were collected, and the concentration of DNA in the solution was measured. The experiment result demonstrated that 1.5D, a pure DNA hydrogel, was highly stable in PBS with a loss of only 1.6% of the total DNA in 21 days **(Figure 3A)**. Similarly, 1.5D2S and 1.5D4S also demonstrated very high stability in PBS with no significant difference in percentage degradation in comparison to 1.5D. These results demonstrate that DNA hydrogels are highly stable in a PBS environment, and even though PBS contains salt, which causes DNA network expansion, due to the absence of the DNase enzyme has a minor impact on DNA degradation. Therefore, we next tested the stability of DNA-Silk hydrogel in DMEM cell culture media supplemented with fetal bovine serum, which contains both salts and nucleases. Similar to the degradation study with PBS, we incubated DNA-Silk fibroin hydrogel in DMEM complete media and collected the media at specific intervals and quantified the total DNA concentration. As expected, pure DNA hydrogel (1.5D) underwent significant DNA degradation in just 1 day **(Figure 3B)**. After 24 hrs, 1.5D hydrogel underwent a 31% DNA weight loss, with almost 91% DNA weight loss at the end of day 3. At the end of day 4, a complete dissolution of the DNA hydrogel occurred. However, both 1.5D2S and 1.5D4S demonstrated significant resistance to DNA degradation. In comparison, 1.5D2S and 1.5D4S hydrogels lost 15.53% and 8.73% of their total DNA at the end of day 4, at which complete dissolution of pure DNA hydrogel occurred. Further, 1.5D2S and 1.5D4S hydrogels were stable even at day 21, with 70% and 55% DNA weight loss. Also, in comparison to 1.5D2S hydrogel, 1.5D4S hydrogels demonstrated higher stability, which can be attributed to the higher concentration of silk fibroin in the hydrogel. These results demonstrate that the introduction of a silk fibroin in a DNA network significantly increases its stability the rate of degradation can be controlled by varying silk concentration. Overall, DNA-Silk fibroin hydrogel overcomes the limited stability limitation of DNA hydrogels and paves the way for the utilisation of DNA-based hybrid hydrogel for applications requiring controllable degradation rates.

**Figure 3:**
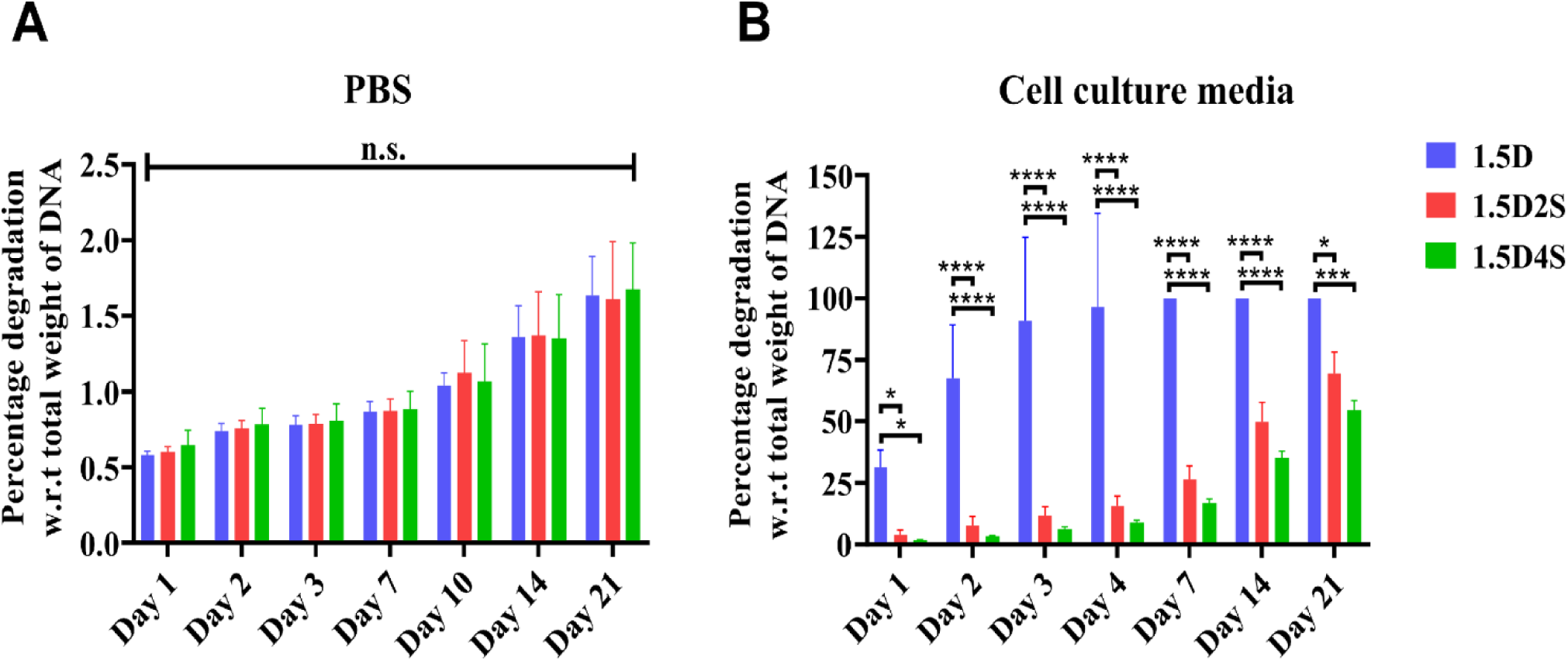
DNA degradation study of DNA-Silk fibroin hydrogels. **(A)** DNA degradation profile of DNA-Silk fibroin hydrogel in PBS and (B) cell culture medium for 21 days. Data presented as mean ± SD. * indicate statistical significance within respective groups. *,***, **** indicate p<0.05, p<0.001, and p<0.0001 respectively, and n.s. indicates no significant difference.

### 2.4 Biocompatibility of DNA-Silk fibroin hydrogel

The cytobiocompatibility of the developed DNA-Silk fibroin hydrogel was evaluated using *in vitro* live/dead staining and MTT cell viability assay. Live/dead fluorescence staining revealed a predominance of green-fluorescent live cells with minimal to no red-fluorescent dead cells after incubation with the hydrogel condition media, indicating negligible cytotoxic effects of both 1.5D2S and 1.5D4S DNA-Silk fibroin hydrogels **(Figure 4A)**. Additionally, in comparison to 1.5D, the pure DNA hydrogel group, cells grown on DNA-Silk fibroin hydrogel, demonstrated an elongated morphology with elongated pseudopodia, suggesting that DNA-Silk fibroin hydrogel provided an environment that promoted cellular attachment and proliferation. Additionally, quantitative evaluation using the MTT assay demonstrated that cell viability in hydrogel condition media-treated groups remained above 84% and 93% after 24 and 48 hrs for 1.5D2S and 1.5D4S, respectively **(Figure 4B)**. However, a significant decrease in cell viability to less than 60% after 48 hrs was observed in the 1.5D pure DNA hydrogel group. This decrease in cell viability can be attributed to the significant amount of degraded DNA present in 1.5D conditioned media, which is known to be involved in reactive oxygen species production and stress response in cells^33,34^. The excellent cytocompatibility observed can be attributed to the hydrogel’s natural component composition and absence of cytotoxic leachates, as well as to its hydrated, porous architecture, which supports nutrient diffusion. Collectively, these findings confirm that the DNA-Silk fibroin hydrogels possess excellent biocompatibility and are appropriate for biological applications requiring direct and prolonged interaction with living cells, including wound healing, tissue engineering, and drug delivery systems. Building upon this excellent biocompatibility profile, subsequent in vitro experiments were conducted to investigate cell-material interactions in greater depth, focusing on proliferation, morphology, and functional behaviour over extended culture periods.

**Figure 4:**
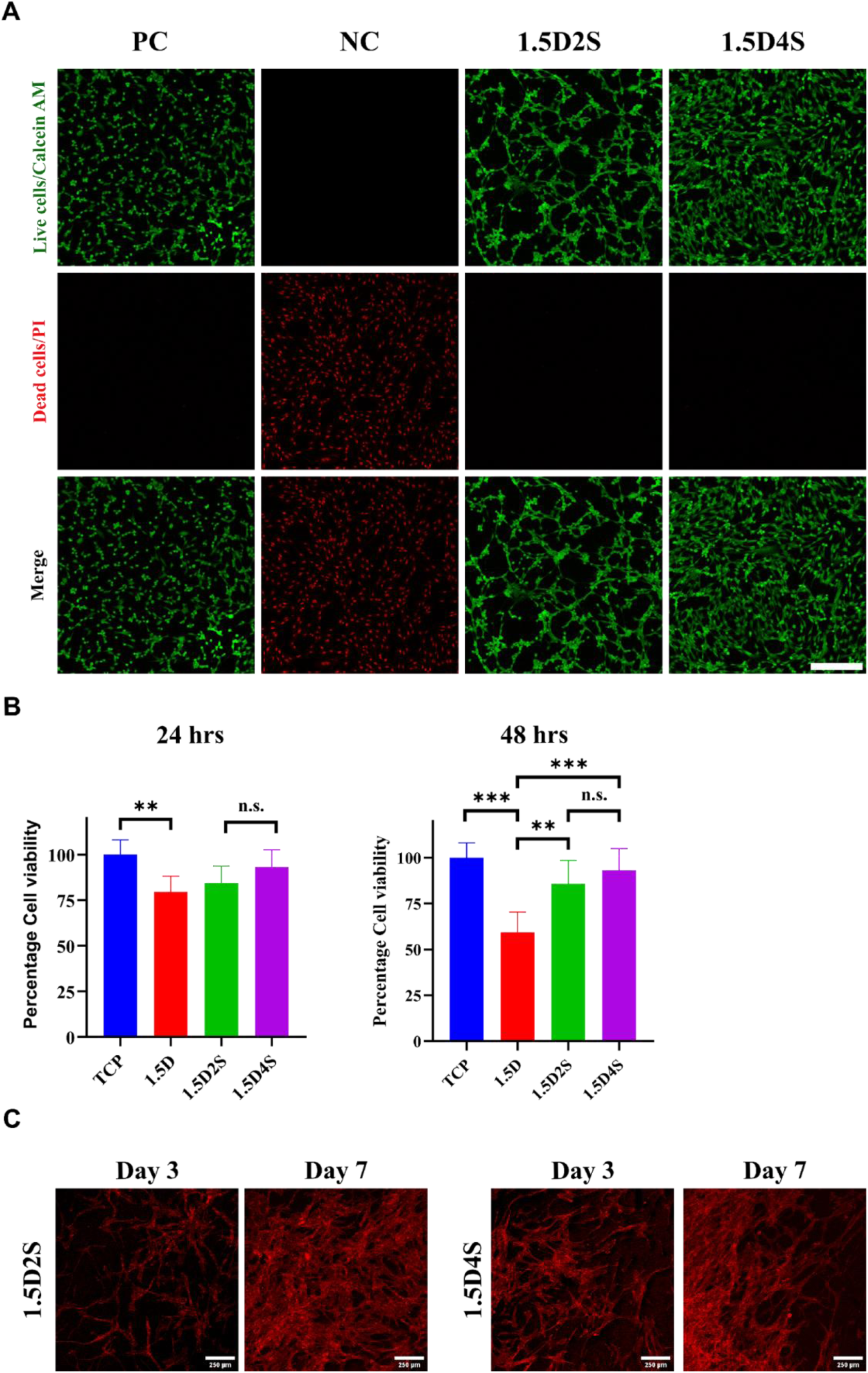
Biocompatibility and in vitro cell culture of DNA-Silk fibroin hydrogel. (A) Live/Dead staining of cells grown on DNA-Silk fibroin hydrogel. Live cells (green) were stained with Calcein AM, and dead cells (red) were stained with Propidium iodide (PI). Cells grown on coverslips were taken as a positive control (PC), and ethanol-killed cells were taken as a negative control (NC). Scale bar-250 µm. (B) Quantitative evaluation of cell viability of cells grown on hydrogels using the MTT assay. Data presented as mean ± SD. * indicate statistical significance within respective groups. **, *** indicate p<0.01 and p<0.001 respectively, and n.s. indicates no significant difference. TCP-Tissue Culture Plate. (C) In vitro cell culture of RPE1 cells on DNA-Silk fibroin hydrogels. Representative confocal microscope images for visualization of the growth pattern of RPE cells cultured on DNA-Silk fibroin by phalloidin staining (red) of F-actin at day 3 and day 7. Scale bar - 250 µm.

### 2.5 In vitro cell culture of DNA-Silk fibroin hydrogel

Building upon the excellent cytocompatibility profile of DNA-Silk fibroin hydrogel, we next evaluated the capacity of the hydrogel to support cell proliferation using human retinal pigment epithelial (RPE1) cells cultured directly on the hydrogel surface. Cells were seeded at a density of 25,000 cells/cm² and maintained under standard culture conditions (37 °C, 5% CO₂) for 3 days and 7 days, with media change every 2 days. To visualise cytoskeletal organisation, cells were fixed and stained with phalloidin to label filamentous actin (F-actin), followed by imaging using confocal microscopy. We observed a dense and interconnected network of RPE1 cells on both 1.5D2S and 1.5D4S hydrogel, indicating that DNA-Silk hydrogel promoted extensive actin cytoskeleton organisation, with cells’ pseudopodia spanning across the hydrogel surface and forming a network-like pattern **(Figure 4C)**. The confocal micrograph further suggests that RPE1 cells strongly adhered to the hydrogel, spread extensively, and established strong cell-hydrogel interactions. The presence of continuous and extensive actin-rich regions further implies robust cytoskeletal integrity and strong focal adhesion complexes, which are critical for cell proliferation and function. Overall, DNA-Silk fibroin hydrogel acts as a promising scaffold material for cell culture applications where the hydrogel can you used for maintaining, proliferating, and potentially differentiating cells.

### 2.6 MSCs Differentiation potential of DNA-Silk fibroin hydrogel

After observing the potential of DNA-Silk fibroin hydrogel as a promising scaffold material for cell culture applications, we investigated whether DNA-Silk fibroin hydrogel can be used as a matrix to support the growth, proliferation and differentiation of mesenchymal stem cells. Thus, we seeded DNA-Silk fibroin hydrogels, which were kept at 37 °C for 3 days with goat derived IFP-MSCs. After 14 days, the evaluation of the differentiation potential of DNA-Silk fibroin hydrogel was conducted by staining the IFP-MSCs using oil red O (adipogenesis), alcian blue (chondrogenesis), and alkaline phosphatase (osteogenesis) staining. For the adipogenic differentiation, we observed that 1.5D2S and 1.5D4S hydrogel demonstrated noticeably reduced lipid droplet density in comparison to the positive control **(Figure 5A)**. This decrease in lipid droplet density can be attributed to increased stiffness and crosslinking within these hydrogels, which may not be suitable for adipogenic differentiation of IFP-MSCs. For chondrogenic differentiation, alcian blue staining revealed enhanced glycosaminoglycan (GAG) deposition in both 1.5D2S and 1.5D4S hydrogels compared to the positive control **(Figure 5B)**. The deep blue colour staining around the cells indicates that the 1.5D2S and 1.5D4S hydrogel supports chondrogenic differentiation of MSCs and promotes chondrogenic signalling pathways and extracellular matrix synthesis. However, no significant difference IN alcian blue staining intensity was observed between the 1.5D2S and 1.5D4S group. These results indicate that the difference in mechanical strength between the 1.5D2S and 1.5D4S hydrogel did not influence the degree of chondrogenic differentiation between the hydrogel groups. However, the influence of the difference in mechanical strength between 1.5D2S and 1.5D4S was observed for the differentiation potential of MSCs towards osteogenic lineage. First, both 1.5D2S and 1.5D4S hydrogels demonstrated higher alkaline phosphatase activity in comparison to the positive control, indicating that MSCs favour DNA-Silk fibroin hydrogel as a matrix in comparison to the tissue culture well plate **(Figure 5C)**. Further, between 1.5D2S and 1.5D4S, the 1.5D4S DNA-Silk fibroin hydrogels resulted in higher alkaline phosphatase staining in comparison to the 1.5D2S hydrogel, suggesting that the higher crosslinking density and increased mechanical stiffness of 1.5D4S likely provided osteoinductive cues to MSCs, resulting in higher differentiation of MSCs towards osteogenic lineage. Further, the osteogenic potential of DNA-Silk hydrogel can be further attributed to the presence of DNA in the hydrogel, which is known to promote bone mineralisation by acting as a nucleation site for calcium phosphate. Overall, these results demonstrate the potential of DNA-Silk hydrogel for cartilage as well as bone tissue engineering. The 1.5D4S DNA-Silk hydrogel, with its superior osteogenic induction capability, may serve as a scaffold for bone regeneration, while 1.5D2S could be utilised for cartilage repair and joint tissue engineering. Thus, the DNA-Silk fibroin hydrogels can act as potential biomaterials for designing mechanically adaptive hydrogels capable of directing stem cell fate for tissue engineering applications.

**Figure 5:**
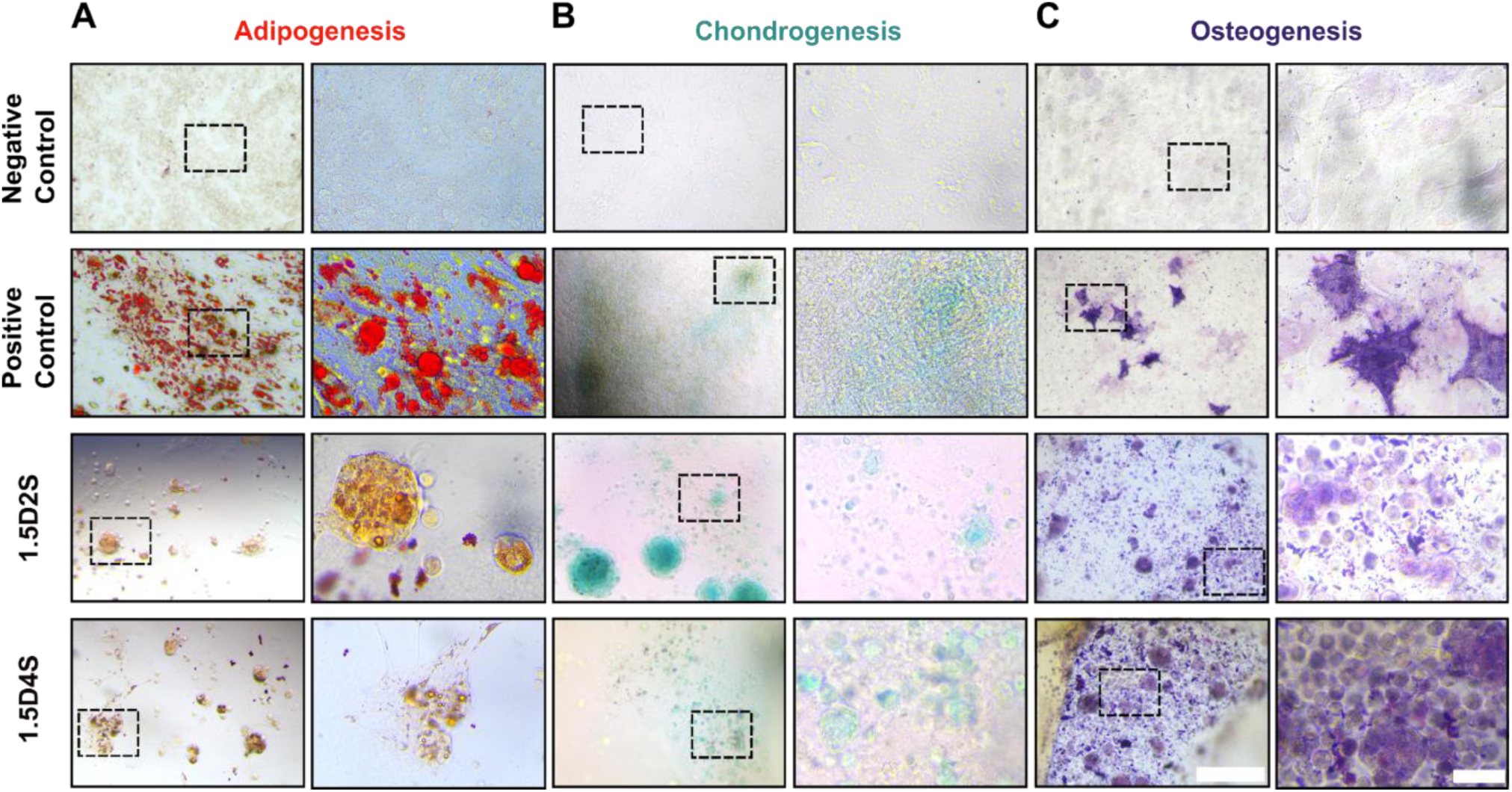
Differentiation potential of DNA-Silk fibroin hydrogel. (A) Oil red O staining of IFP-MSCs. (B) Alcian blue staining of IFP-MSCs. (C) Alkaline phosphatase staining of IFP-MSCs. The right panel shows the magnified images of the area marked by the dashed box (scale bar-50 µm). The Left panel has a scale bar of 250 µm.

### 2.7 In vitro hemostasis and hemocompatibility of DNA-Silk fibroin hydrogel

In recent years, extensive research has been done towards developing materials that can not only be utilised for wound healing applications but can also be applied at the wound site to prevent traumatic bleeding. Hydrogels have emerged as a potential material to prevent bleeding and simultaneously promote wound healing, as they can absorb a significant amount of blood at the wound site, preventing blood loss, and can take the shape of the wound site to promote wound healing^35–37^. Among different biomaterials, DNA hydrogels offer a significant advantage in comparison to other polymers, as it can mimic neutrophil extracellular traps (NETs), which is a network of DNA and proteins released by neutrophils in response to injury to trap pathogens as well as platelets to promote blood coagulation^38,39^. Hence, DNA hydrogel can act as an excellent material and can be used as a dressing material to prevent blood loss and promote wound healing. Thus, we next investigated whether DNA-Silk fibroin hydrogel can be utilised as a hemostatic biomaterial that can coagulate blood and simultaneously promote wound healing.

We first investigated whether DNA-Silk fibroin hydrogels are hemocompatible or not. To test the hemocompatibility of DNA-Silk hydrogel, we performed the hemolysis assay, and the results showed that both 1.5D2S and 1.5D4S hydrogels are highly hemocompatible, with 1.5D2S demonstrating negligible hemolysis and 1.5D4S with a hemolysis percentage of 1.5% **(Figure 6A)**. As the hemolysis ratio is below 5% (ASTM standard), the developed hydrogels are hemocompatible and thus can be used for biological applications. We next evaluated the blood-clotting ability of DNA-Silk fibroin hydrogel. We performed in vitro hemostasis test and observed that both 1.5D2S and 1.5D4S hydrogels resulted in blood coagulation within 10 mins, as indicated by the absence of blood at the bottom of the centrifuge tube **(Figure 6B)**. The result demonstrates that DNA-Silk hydrogel has the ability to absorb and coagulate blood. To further test the hemostatic ability of DNA-Silk fibroin hydrogel, we measured the blood clotting index of the hydrogels. We observed that in comparison to the control group, where no hydrogel was there, both 1.5D2S and 1.5D4S hydrogel absorbed and coagulated the blood, as observed by the clear solution in the petri dish in comparison to the control **(Figure 6C)**. We further quantified the blood clotting index by measuring the absorbance of haemoglobin in the petri dish solution and observed that both 1.5D2S and 1.5D4S hydrogel resulted in significantly lower BCI of 28.36 ± 7.187 % and 19.02 ± 6.955 %, respectively, in comparison to the control. Overall, these results demonstrate the hemocompatibility and blood clotting ability of DNA-Silk fibroin hydrogel and demonstrate that the DNA-Silk hydrogel can be used as a hemostatic material to stop bleeding during injury and opens the potential of DNA-Silk hydrogel as a dressing material for wound healing applications. Thus, we next evaluated the wound healing ability of DNA-Silk fibroin hydrogel.

**Figure 6:**
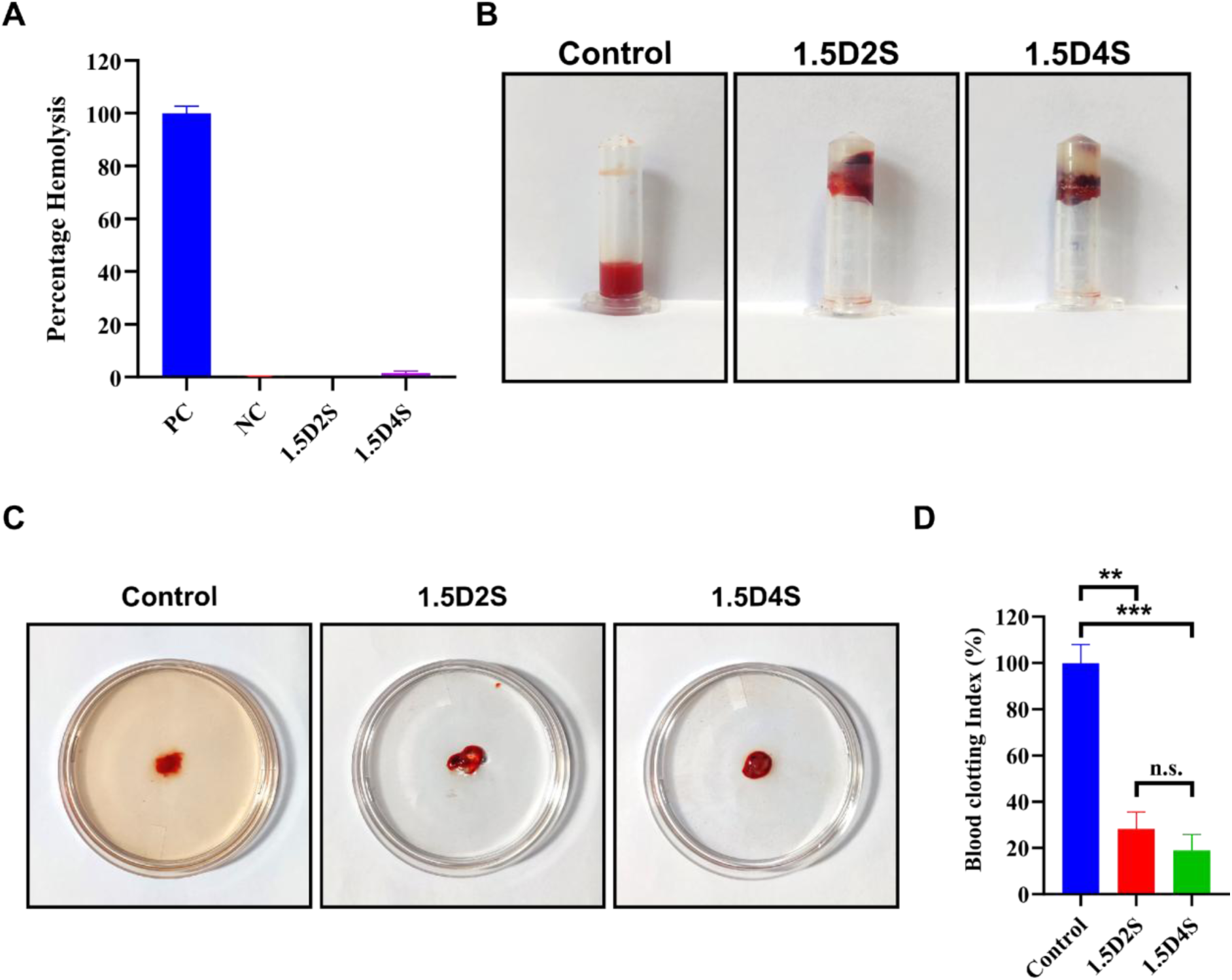
In vitro hemostasis and hemocompatibility of DNA-Silk fibroin hydrogel. (A) In vitro hemolysis assay of DNA-Silk fibroin hydrogel. PC-Positive Control, NC-Negative Control. (B) In vitro hemostatic assay of DNA-Silk hydrogel. (C) Evaluation of the blood clotting ability of DNA-Silk hydrogel. (D) Blood clotting index assay of DNA-Sil fibroin hydrogel. **, *** indicate p<0.01 and p<0.001 respectively, and n.s. indicates no significant difference.

### 2.8 In vivo wound healing study of DNA-Silk fibroin hydrogel

Due to superior hemocompatibility and hemostatic ability, we next investigated the wound healing ability of DNA-Silk fibroin hydrogel in treating the full-thickness skin defect in BALB/c mice. The therapeutic efficacy of the developed hydrogel was monitored by measuring the reduction in wound area over the course of 11 days. The wound size area was measured at day 3, day 6, day 9 and day 11. By calculating the percentage wound closure, we quantified the healing ability of the DNA-Silk fibroin hydrogels. We observed that in comparison to both the negative and positive controls, DNA-Silk hydrogel resulted in faster wound healing as indicated by decreased wound area in comparison to the control group **(Figure 7A and 7B)**. Between 1.5D2S and 1.5D4S hydrogel, 1.5D4S hydrogel demonstrated a higher reduction in wound area. Quantification of wound area and calculation of percentage extent of healing further corroborated these results with 1.5D2S and 1.5D4S hydrogel, resulting in significant wound contraction of 40.09% and 44.19% at the end of day 3, respectively **(Figure 7C)**. In comparison, the negative control group (untreated mice) and the positive control group (silver nitrate gel-treated mice) resulted in wound contraction of 9.61% and 14.49%, respectively. Similar results were obtained at the end of day 6, with 1.5D4S hydrogel resulting in significantly higher wound closure in comparison to the control groups. The extent of wound healing of the DNA-Silk hydrogel continued to be higher than the control group at day 9 and day 11, also. Overall, DNA-Silk hydrogels, especially 1.5D4S, demonstrated accelerated wound healing in comparison to the control group based on the rate of wound closure. To further investigate the effect of DNA-Silk hydrogels on wound healing at the tissue level, histological evaluation of the wound site skin was performed. Hematoxylin and Eosin (H&E) staining of the 1.5D4S hydrogel-treated wound tissue section demonstrated that 1.5D4S resulted in significant wound healing with morphology similar to that of normal skin tissue **(Figure 7D)**. Notably. new hair follicles, new epithelium, granular tissue and blood vessels were observed in both 1.5D2S and 1.5D4S hydrogel groups. Additionally, Sirius red staining for collagen demonstrated a significantly higher amount of collagen deposition in both 1.5D2S and 1.5D4S groups in comparison to the control groups, indicating that treatment of the wound with DNA-Silk fibroin hydrogel resulted in complete dermal tissue regeneration. Overall, DNA-Silk fibroin hydrogel resulted in accelerated wound healing with regeneration of all skin layers and components.

**Figure 7:**
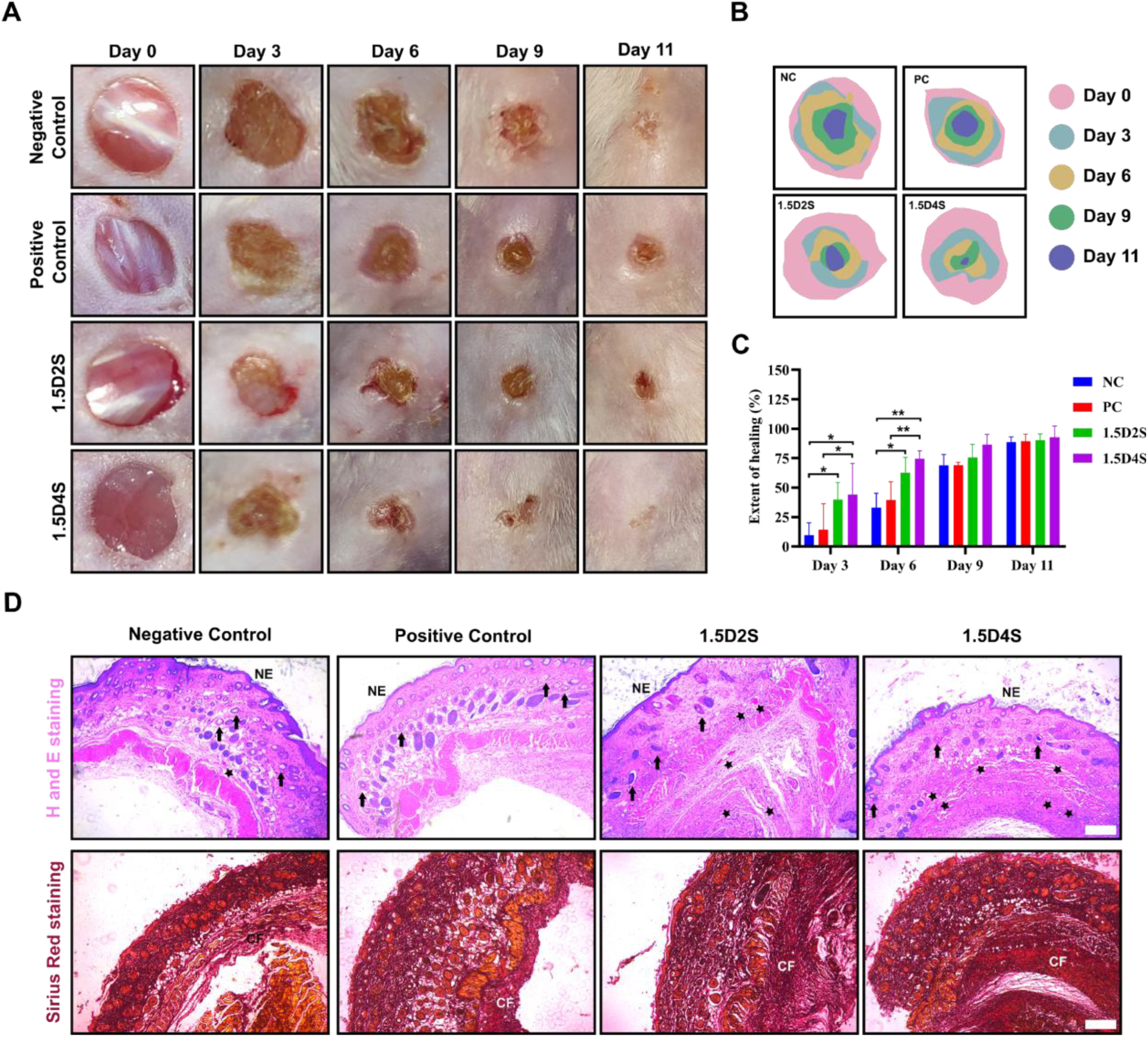
In vivo wound healing study of DNA-Silk fibroin hydrogel. (A) Wound images of mice with a full skin defect at different timepoints. (B) Trace micrograph of wound healing on different days. (C) Quantification of the extent of wound healing based on wound size at different time points. NC-Negative Control, PC-Positive Control. *, ** indicate p<0.05 and p<0.01 respectively. (D) Histopathological staining of the wound site skin using H&E and Sirius red staining. Star indicates new blood vessels, arrow points towards new hair follicles, NE-new epithelium, CF-Collagen fibre. Scale bar-250 µm.

## 3. Conclusions

In this study, we successfully developed a DNA-silk fibroin hybrid hydrogel that addresses the key limitations of conventional DNA-based hydrogels, including high cost, poor mechanical strength, limited stability, and complex synthesis. By utilising salmon-derived DNA as a cost-effective and sustainable source, and silk fibroin as a mechanically robust and biocompatible scaffold, we achieved a hybrid material with superior mechanical properties, enhanced structural stability, and controllable degradability. The synergistic interaction between DNA and silk fibroin resulted in a dynamic mechanical reinforcement of the hydrogel with an entangled network that not only preserved the functional advantages of DNA but also significantly improved its durability and applicability. Beyond its physicochemical benefits, the hybrid hydrogel exhibited excellent biocompatibility, supporting robust cell attachment, spreading, and proliferation in vitro. Additionally, DNA-Silk fibroin hydrogels supported both chondrogenic and osteogenic lineage commitment, highlighting its suitability for cartilage and bone tissue engineering applications. In addition, the presence of DNA, a key component of the neutrophil extracellular trap (NET) matrix, endowed the hydrogel with inherent hemostatic potential, facilitating rapid clot formation and indicating its promise as a wound-hemostatic adjuvant. Finally, in vivo evaluation using a full skin defect mice model confirmed that the hybrid hydrogel significantly accelerated wound closure and tissue regeneration, highlighting its potential as an effective wound-healing material. Together, these findings establish the DNA-Silk fibroin hydrogel as a multifunctional biomaterial platform capable of promoting stem cell differentiation, achieving rapid hemostasis, and facilitating tissue repair. Moving forward, its tunable mechanical and biochemical properties can be leveraged for broader biomedical applications, including regenerative medicine, implantable scaffolds, advanced wound dressings, and drug delivery systems.

## 4. Materials and methods

### 4.1 Materials

DNA sodium salt, lithium bromide, calcium chloride, propidium iodide, insulin, transferrin, 4-(2-hydroxyethyl)-1-piperazineethanesulfonic acid (HEPES), β-glycerophosphate, 2-Phospho-L-ascorbic acid trisodium salt, sodium selenite, indomethacin and sodium carbonate were purchased from Sigma-Aldrich. Poly(ethylene glycol) (MW:20000 Da), proline, Vitamin D3 and dexamethasone were procured from Himedia. Ethanol was procured from CSS Fine Chemicals. Calcein AM and MTT (3-(4,5-dimethylthiazol-2-yl)-2,5-diphenyltetrazolium Bromide) were procured from Invitrogen, ThermoFisher Scientific. Dulbecco’s Modified Eagle Medium (DMEM), Penicillin-Streptomycin (10,000 U/mL), and Fetal Bovine Serum (FBS) were purchased from Gibco, ThermoFisher Scientific. Bombyx *mori* cocoons were purchased from a local silk mill vendor.

### 4.2 Extraction of Silk fibroin from B. *mori* cocoons

Extraction of silk fibroin from B. *mori* cocoons was done following a previously published protocol^40^. Briefly, 5 gm of B. *mori* cocoons were cut into small pieces (5 mm x 5 mm) and boiled in 2 L of 0.02 M sodium carbonate solution for 30 min. After boiling, the obtained silk fibres were rinsed three times in 1 L of ultrapure water for 20 min. The silk fibroin fibres were kept for drying in a fume hood overnight. Then 1 gm of silk fibroin fibres were dissolved in 10 ml of 9.3 M Lithium bromide solution for 4hrs at 60 °C. Upon complete dissolution, the silk fibroin salt solution was dialysed (MWCO: 12 KDa) against 2 L of ultrapure water for 3 days with repeated water changes every 6 hrs. After dialysis, the silk fibroin aqueous solution was centrifuged twice at 12700 g for 20 minutes at 4°C. To concentrate the silk fibroin solution, the solution was dialyzed against a 10 w/v% poly(ethylene glycol) solution for 12 hrs. To determine the concentration of silk fibroin solution, the dry weight analysis method was used. Briefly, 0.5 ml of silk fibroin solution was added to a weigh boat and allowed to dry at 60 °C. After drying silk weight was divided by 0.5 ml to determine the concentration of silk fibroin in solution.

### 4.3 Preparation of DNA-Silk Hybrid Hydrogel

To prepare the DNA-Silk fibroin hybrid hydrogel, a DNA sodium salt solution was mixed with Silk fibroin (SF) solution in different weight percentage ratios **(Table 1)**. For all the experiments, a 1.5 w/v% concentration of DNA was maintained, and the ratio of SF solution was varied. 1.5D was taken as the DNA control, and 2S and 4S were taken as the silk control. After mixing, the DNA-silk solution was heated to 93°C for 7 min at 1000 rpm, and ethanol was added during stirring to induce rapid silk gelation (20% ethanol concentration with respect to the reaction volume). After stirring, the DNA-Silk solution was kept at 37°C in an environment with more than 90% relative humidity.

### 4.4 Physical and chemical characterisation

The morphological analysis of DNA-Silk hydrogel was carried out by scanning electron microscopy (SEM). For SEM analysis, the DNA-Silk hydrogels were first freeze-dried and subsequently sputter-coated with gold and observed at 5 kV acceleration voltage. For mechanical characterisation, rheological analysis of the DNA-Silk hydrogel was performed using a rheometer. Frequency sweeps were taken at the constant shear strain of 1% with a frequency range from 1 Hz to 100 Hz at 25°C. An amplitude sweep was performed at a constant 1% shear strain at 25°C. FTIR analysis was conducted in the range of 400 cm^-^ to 4000 cm^−^. To observe the β-sheet content of the DNA-Silk hydrogel, Fourier self-deconvolution was performed on the region corresponding to β-sheets.

### 4.5 DNA Degradation study of DNA-Silk hybrid Hydrogel

To study the degradation of DNA-Silk hybrid hydrogel, the hydrogels (5-day incubated) was incubated in 500 μL of PBS or DMEM media with 10% FBS at 37°C for different numbers of days (1,2,3,4,7,10,14 days). After the respective time point, the supernatant was collected, and the amount of DNA was measured spectroscopically using a spectrophotometer (NanoDrop 2000/2000c).

### 4.6 In vitro cell culture

Human retinal pigment epithelial-1 (RPE-1) cells and Goat infrapatellar fat pad-derived mesenchymal stem cells (IFP-MSCs) were used for *in vitro* cell culture studies. Goat IFP-MSCs were harvested from the goat knee following a standard published protocol^41^. RPE-1 cells were grown in DMEM high-glucose media supplemented with 10% FBS and 1% penicillin-streptomycin and were used for biocompatibility tests. Goat IFP-MSCs were used for differentiation studies on DNA-Silk hybrid hydrogel. Goat IFP-MSCs were maintained in DMEM expansion media (DMEM Low glucose, 10% FBS, 100 U/mL penicillin-streptomycin, and 2.5 μg/mL amphotericin). For all *in vitro* cell culture experiments, a temperature of 37°C and 5% CO_2_ was maintained inside the incubator.

### 4.7 Biocompatibility tests

In vitro cytocompatibility of DNA-Silk hybrid hydrogel was evaluated using MTT and Live-Dead assay. For the MTT assay, RPE-1 cells were treated with DNA-Silk fibroin hybrid hydrogel (5-day incubated) conditioned media. The DNA-Silk hybrid hydrogel conditioned media was obtained by incubating the hydrogels with 500 μL DMEM high-glucose media for 24 and 48 hrs at 37°C and 5% CO_2_. Initially, RPE-1 cells were seeded in a 96-well plate with a seeding density of 25000 cells/ml for 24 hrs. After 24 hrs, RPE-1 cells were treated with conditioned media for 24 hrs and 48 hrs. RPE-1 cells incubated with pure DMEM media was taken as the control. After 24 or 48 hrs, conditioned media was removed, and cells were incubated with DMEM serum-free media containing 0.5mg/ml MTT dye (3-(4,5-Dimethylthiazol-2-yl)-2,5-diphenyltetrazolium Bromide) for 4 hrs at 37°C. After 4 hrs, MTT dye was removed, and absorbance was measured at 370 nm in DMSO to quantify cell viability. For the Live/Dead assay, RPE-1 cells were incubated with DNA-Silk hybrid hydrogel conditioned media for 3 days. After 3 days, cells were stained with 1.5 µg/ml of Calcein AM and propidium iodide for 10 min at 37 °C in a cell culture incubator at 5% CO_2_, in the absence of light. After staining, the cells were washed with 1X PBS and were imaged using a confocal laser scanning microscope. For the negative control, the cells were killed using ethanol for 10 minutes, followed by staining with Calcein AM and propidium iodide.

### 4.8 Differentiation of Goat Infrapatellar Fat Pad-Derived Mesenchymal Stem Cells

Goat IFP-MSCs were used for differentiation studies on DNA-Silk hybrid hydrogel. For all in vitro cell culture experiments, a temperature of 37°C and 5% CO2 was maintained inside the incubator. For the differentiation study, IFP-MSCs were seeded on hydrogels incubated a cell culture well plate for 3 days. For adipogenesis, chondrogenesis and osteogenesis, 25000 cells/cm^2^, 50000 cells/cm^2^ and 25000 cells/cm^2^ were seeded in a cell culture plate, respectively. The cells were grown in the respective induction media (**Adipogenesis induction media**: DMEM expansion media supplemented with 25 mM D(+) glucose, 1 μM dexamethasone, 10 μg/mL insulin, and 100 μM indomethacin; **Chondrogenic induction media**: DMEM expansion media supplemented with 10 mM 4-(2-hydroxyethyl)-1-piperazineethanesulfonic acid(HEPES), 1 mM proline, 100 nM dexamethasone, 10 µg/mL insulin, 5.5 µg/mL transferrin, 6.7 ng/mL sodium selenite, 1.25 mg/mL bovine serum albumin (BSA), 50 μg/mL 2-Phospho-L-ascorbic acid trisodium salt; **Osteogenic induction media**: DMEM expansion media supplemented with 50 μg/mL 2-Phospho-L-ascorbic acid trisodium salt, 25 mM D (+)-glucose, 10 mM β-glycerophosphate, 10 nM dexamethasone and 10 nM vitamin D3) with media changes every 3 days. For the negative control (uninduced), cells were grown in a tissue cell culture plate in DMEM expansion media supplemented with 25 mM D (+) glucose. For positive control, cells were grown in a tissue cell culture plate supplemented with the respective induction media. The cell culture was continued till day 14, after which histological staining of IFP-MSCs was performed to observe cellular differentiation. For adipogenesis, oil droplets were visualised using oil red staining as per a previously reported standard histology protocol^41^. Similarly, for chondrogenesis, the proteoglycan content in the matrix and for osteogenesis, the alkaline phosphatase activity were evaluated using alcian blue and alkaline phosphatase staining, respectively, following a previously reported standard histology protocol^41^.

### 4.9 Hemocompatibility test

The hemocompatibility test of the DNA-Silk hydrogel was assessed by determining the percentage hemolysis of red blood cells (RBCs). Briefly, fresh blood was collected from mice and centrifuged at 1500 g for 10 min to collect red blood cells. The collected red blood cell pellet was washed with 1X PBS three times and resuspended in 1X PBS in a 2% ratio(v/v). 10 mg of DNA-Silk fibroin hydrogel (3-day incubated) was then incubated with 500 μL of 2% RBCs solution and shaken at 37° C for 1 h at 200 rpm. After 1 h, the samples were centrifuged at 1500 g for 10 min to remove RBCs, and the supernatant was collected for UV absorbance. The percentage hemolysis was measured by determining the amount of haemoglobin in the supernatant by UV absorbance at 540 nm. The percentage hemolysis was calculated following the equation: Percentage hemolysis (%) = (A_s_/A_o_) x 100, where A_s_ is the absorbance of the sample and A_o_ is the absorbance of the positive control. For the test, the RBCs solution treated with 0.1% Triton-X solution was taken as a positive control, and the RBCs suspended in 1X PBS were taken as a negative control.

### 4.10 In vitro hemostatic performance of DNA-Silk hydrogel

To investigate the hemostatic performance of the DNA-Silk fibroin hydrogel, 500 μL of DNA-Silk hydrogel was added to 2 mL centrifuge tubes. The hydrogels were incubated at 37°C for 10 minutes, and subsequently, 200 μL anticoagulated mice blood was added to the hydrogels. The hydrogels were incubated with blood for 5 min, followed by the addition of 50 μL of 25 mM calcium chloride for 5 min. The centrifuge tubes were tilted every 30 seconds to see whether the blood had coagulated or not. After coagulation, the centrifuge tubes were inverted and photographed.

### 4.11 Blood Clotting Index test

The BCI test was performed to test the hemostatic ability of DNA-Silk fibroin hydrogel. Briefly, the DNA-Silk hydrogels were pre-heated to 37°C for 10 mins, and 200 μL of DNA-Silk hydrogel was then placed in the centre of a 90 mm petri dish. Then, 50 μL of freshly collected mice blood was added on top of the hydrogels and kept for 5 min. After 5 minutes, 10 μL of 0.2 M calcium chloride solution was added and kept for another 5 minutes. Then, 10 mL of distilled water was added to the petri dish, and the petri dish was shaken slowly, not to disturb the hydrogel, to collect RBCs not trapped in the hydrogel. After 5 min, the solution containing free RBCs was collected, and the UV absorbance of the collected solution was measured at 540 nm. The blood clotting Index (BCI) was calculated following the equation: BCI (%) = (A_s_/A_o_) x 100, where A_s_ is the absorbance of the sample and A_o_ is the absorbance of the control group. For the control group, 50 μL of freshly collected mice blood was added directly onto the petri dish without any hydrogel, followed by the addition of the calcium chloride.

### 4.12 In vivo wound healing

The wound healing studies were performed on animal models of male BALB/c mice weighing 18-23 grams, and they were procured from Zydus Research Centre (ZRC), Ahmedabad-Gujarat. The animals were housed in polypropylene cages with a 12-hour light-dark cycle at a temperature of 22°C. The in-vivo wound healing studies were performed by creating a full-thickness skin defect on the mice following a standard protocol, adhering to ethical and biosafety guidelines. The study was conducted after approval from the Institutional Animal Ethics Committee (IAEC) of Nirma University in Ahmedabad, Gujarat: ISNU/PHD/39/2025/33. The experiments were conducted following the authorised protocol with careful safety and well-being of animals. The mice were divided into 4 groups 3 mice in each group: Negative control (no treatment), Positive control (treatment with commercially available 0.2% w/w silver nitrate gel (Silverex)), 1.5D2S, and 1.5D4S. For in vivo studies, DNA-Silk fibroin hydrogels, which were kept at 37°C in 90% humidity for 3 days, were used. First, the hair on the animal’s back was shaved, and then the mice were mildly anaesthetised with diethyl ether. A circular wound of 5 mm was created on the mice back using a biopsy punch, and then the hydrogels were applied to the wound site. The wound site was then subsequently covered with a dressing to prevent infection. The dressing was changed, and the hydrogels were applied every 3 days. The wound images were taken during every dressing change (day 3, day 6, day 9, and day 11), and the percentage wound closure was calculated following the equation:

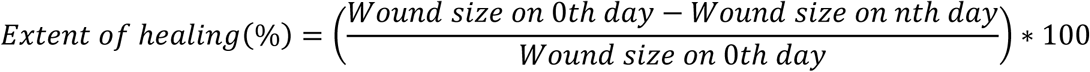

After 11 days, when complete wound closure was observed, the mice were euthanised. The wound site skin was collected, and histopathological studies were conducted.

### 4.13 Histopathological study

The skin specimens collected from the mice’s wound site on Day 11 were fixed in 4% formaldehyde for 24 h. The fixed sections were then embedded in paraffin and sectioned using a microtome to get 5 µm tissue sections. To observe the morphology, tissue regeneration, and collagen distribution after wound healing, the tissue sections were stained using Hematoxylin and Eosin (H&E) and Sirius red staining following standard protocols^41^.

### 4.14 Statistical Analysis

All experiments were independently conducted at least three times, and data are expressed as mean ± standard deviation. Representative results from triplicate experiments are presented. Statistical analyses were performed using GraphPad Prism version 8, employing one-way analysis of variance (ANOVA) and unpaired t-tests, followed by Tukey’s post hoc test or Welch’s correction where appropriate. A p-value < 0.05 was considered statistically significant.

## Acknowledgments

All the members of the D.B. research group are acknowledged for providing constructive comments and a critical review of the manuscript. The infrastructural and financial support from the Indian Institute of Technology Gandhinagar is gratefully acknowledged. N.S., B.P., and A.S. acknowledge the Ministry of Education, Government of India, and A.S. also acknowledges the Prime Minister’s Research Fellowship. Field emission scanning electron microscopy and confocal microscopy analyses were performed with the help of the Central Instrumentation Facility, IITGN. The author also acknowledges S.S. and Nirma University, Ahmedabad, Gujarat, for providing the facility and resources for in vivo experiments. D.B. acknowledges the Science and Engineering Research Board (SERB) and Government of India for the Core Research Grant, IITGN for the start-up grant, and GUJCOST-DST, GSBTM, and STARS-MoES for providing additional research funding. D.B. is also recognised as a member of the Indian National Young Academy of Sciences

## Authors contributions

Conceptualization: NS and DB; Methodology: NS, BP, AJ, DG, AS, RS, VK, SS and DB; Formal analysis and investigation: NS, AJ and DG; Validation: NS, BP, VK and DB; Writing - original draft preparation: NS; Writing - review and editing: NS, AJ, DG, SS and DB; Funding acquisition: DB; Resources: SS and DB; Supervision: DB; Project administration: DB.

## Data availability

All data generated or analysed during this study are included in this published article.

## References

1. Ma, W. et al. The biological applications of DNA nanomaterials: current challenges and future directions. Signal Transduct. Target. Ther. 6, 351 (2021).

2. Singh, N., Singh, A., Dhanka, M. & Bhatia, D. DNA functionalized programmable hybrid biomaterials for targeted multiplexed applications. J. Mater. Chem. B 12, 7267–7291 (2024).

3. Kansara, K., Mansuri, A., Kumar, A. & Bhatia, D. DNA Nano-Biomaterials Based Futuristic Technologies for Tissue Engineering and Regenerative Therapeutics. Small 21, 2504361 (2025).

4. Morya, V., Walia, S., Mandal, B. B., Ghoroi, C. & Bhatia, D. Functional DNA Based Hydrogels: Development, Properties and Biological Applications. ACS Biomater. Sci. Eng. 6, 6021–6035 (2020).

5. Singh, N. et al. DNA-based Precision Tools to Probe and Program Mechanobiology and Organ Engineering. Small 21, 2410440 (2025).

6. Zhan, P. et al. Recent Advances in DNA Origami-Engineered Nanomaterials and Applications. Chem. Rev. 123, 3976–4050 (2023).

7. Seeman, N. C. & Sleiman, H. F. DNA nanotechnology. Nat. Rev. Mater. 3, 17068 (2017).

8. Pang, C., T. Karlinsey, B. & T. Woolley, A. From biology to circuitry: a review of DNA and other biomaterials as templates for nanoelectronic systems. Chem. Commun. 61, 11551– 11566 (2025).

9. Ma, N. et al. Environment-Resistant DNA Origami Crystals Bridged by Rigid DNA Rods with Adjustable Unit Cells. Nano Lett. 21, 3581–3587 (2021).

10. Agarwal, S., Klocke, M. A., Pungchai, P. E. & Franco, E. Dynamic self-assembly of compartmentalized DNA nanotubes. Nat. Commun. 12, 3557 (2021).

11. Ullah, S. et al. Electrospun composite nanofibers of deoxyribonucleic acid and polylactic acid for skincare applications. J. Biomed. Mater. Res. A 111, 1798–1807 (2023).

12. Buchberger, A. et al. Bioactive Fibronectin-III10–DNA Origami Nanofibers Promote Cell Adhesion and Spreading. ACS Appl. Bio Mater. 5, 4625–4634 (2022).

13. Wu, R. et al. DNA hydrogels and their derivatives in biomedical engineering applications. J. Nanobiotechnology 22, 518 (2024).

14. Singh, A. & Bhatia, D. Chapter 16 - DNA hydrogels: Principles, synthesis, characterization and applications to cell biology. in Methods in Cell Biology (ed. Shukla, A. K.) vol. 169 323–346 (Academic Press, 2022).

15. Zhou, L. et al. Functional DNA-based hydrogel intelligent materials for biomedical applications. J. Mater. Chem. B 8, 1991–2009 (2020).

16. Jian, X. et al. Development, Preparation, and Biomedical Applications of DNA-Based Hydrogels. Front. Bioeng. Biotechnol. 9, (2021).

17. Tang, J., Ou, J., Zhu, C., Yao, C. & Yang, D. Flash Synthesis of DNA Hydrogel via Supramacromolecular Assembly of DNA Chains and Upconversion Nanoparticles for Cell Engineering. Adv. Funct. Mater. 32, 2107267 (2022).

18. Yao, C., Zhang, R., Tang, J. & Yang, D. Rolling circle amplification (RCA)-based DNA hydrogel. Nat. Protoc. 16, 5460–5483 (2021).

19. Masukawa, M. K., Okuda, Y. & Takinoue, M. Aqueous Triple-Phase System in Microwell Array for Generating Uniform-Sized DNA Hydrogel Particles. Front. Genet. 12, (2021).

20. Shi, Z., Li, Y., Du, X., Liu, D. & Dong, Y. Constructing Stiffness Tunable DNA Hydrogels Based on DNA Modules with Adjustable Rigidity. Nano Lett. 24, 8634–8641 (2024).

21. Lachance-Brais, C. et al. Small Molecule-Templated DNA Hydrogel with Record Stiffness Integrates and Releases DNA Nanostructures and Gene Silencing Nucleic Acids. Adv. Sci. Weinh. Baden-Wurtt. Ger. 10, e2205713 (2023).

22. Tang, J. et al. A DNA/Poly-(L-lysine) Hydrogel with Long Shelf-Time for 3D Cell Culture. Small Methods 8, 2301236 (2024).

23. Yang, B. et al. A Biostable l-DNA Hydrogel with Improved Stability for Biomedical Applications. Angew. Chem. Int. Ed. 61, e202202520 (2022).

24. Menon, D., Singh, R., Joshi, K. B., Gupta, S. & Bhatia, D. Designer, Programmable DNA-peptide hybrid materials with emergent properties to probe and modulate biological systems. ChemBioChem 24, e202200580 (2023).

25. Shin, S. W., Yuk, J. S., Chun, S. H., Lim, Y. T. & Um, S. H. Hybrid material of structural DNA with inorganic compound: synthesis, applications, and perspective. Nano Converg. 7, 2 (2020).

26. Bhunia, B. K., Bandyopadhyay, A., Dey, S. & Mandal, B. B. Silk-hydrogel functionalized with human decellularized Wharton’s jelly extracellular matrix as a minimally invasive injectable hydrogel system for potential nucleus pulposus tissue replacement therapy. Int. J. Biol. Macromol. 127686 (2023) doi:10.1016/j.ijbiomac.2023.127686.

27. Hassan, S. et al. Injectable Self-Oxygenating Cardio-Protective and Tissue Adhesive Silk-Based Hydrogel for Alleviating Ischemia After Mi Injury. Small 20, 2312261 (2024).

28. Farokhi, M. et al. Crosslinking strategies for silk fibroin hydrogels: promising biomedical materials. Biomed. Mater. 16, 022004 (2021).

29. Kim, U.-J. et al. Structure and Properties of Silk Hydrogels. Biomacromolecules 5, 786– 792 (2004).

30. Singh, N., Bhattacharjee, A., Kumar, P. & Katti, D. S. Targeting multiple disease hallmarks using a synergistic disease-modifying drug combination ameliorates osteoarthritis *via* inhibition of senescence and inflammation. Life Sci. 334, 122212 (2023).

31. Zhang, C. et al. Counterintuitive DNA destabilization by monovalent salt at high concentrations due to overcharging. Nat. Commun. 16, 113 (2025).

32. Chandrasekaran, A. R. Nuclease resistance of DNA nanostructures. Nat. Rev. Chem. 5, 225–239 (2021).

33. Silk, E., Zhao, H., Weng, H. & Ma, D. The role of extracellular histone in organ injury. Cell Death Dis. 8, e2812–e2812 (2017).

34. Gottlieb, Y. et al. Neutrophil extracellular traps in pediatric inflammatory bowel disease. Pathol. Int. 68, 517–523 (2018).

35. Ding, R. et al. A Multifunctional Injectable Hydrogel Accelerates Hemostasis and Wound Healing via Synergistic Tissue Adhesion, Antimicrobial Activity, and ROS Scavenging. ACS Appl. Mater. Interfaces 10.1021/acsami.5c14873 (2025) doi:10.1021/acsami.5c14873.

36. Yang, J. et al. Injectable hemostatic hydrogel adhesive with antioxidant, antibacterial and procoagulant properties for hemorrhage wound management. J. Colloid Interface Sci. 673, 395–410 (2024).

37. Wang, P., Pu, Y. & He, B. Natural polysaccharide-based hydrogels for hemostasis and wound healing: A review. Precis. Med. Eng. 2, 100016 (2025).

38. Ye, R. et al. Neutrophil extracellular traps-inspired DNA hydrogel for wound hemostatic adjuvant. Nat. Commun. 15, 5557 (2024).

39. Rada, B. Neutrophil Extracellular Traps. Methods Mol. Biol. Clifton NJ 1982, 517–528 (2019).

40. Rockwood, D. N. et al. Materials Fabrication from Bombyx mori Silk Fibroin. Nat. Protoc. 6, 10.1038/nprot.2011.379 (2011).

41. Mahajan, A., Hazra, S., Arora, A. & Katti, D. S. Isolation, Expansion, and Differentiation of Mesenchymal Stem Cells from the Infrapatellar Fat Pad of the Goat Stifle Joint. J. Vis. Exp. JoVE 10.3791/63617 (2022) doi:10.3791/63617.

